# Hidden multiplicity in the analysis of variance (ANOVA): multiple contrast tests as an alternative

**DOI:** 10.1101/2022.01.15.476452

**Authors:** Ludwig A. Hothorn

## Abstract

In bio-medical studies, the p-values of the F-tests in ANOVA are usually interpreted independently as measures of the significance of the associated factors. This ’hidden multiplicity’ effect increases the false positive rate. Therefore, Cramer et al. (2016) proposed the Bonferroni adjustment of the p-values to control for familywise error rate for the experiment. Here, instead of using F-tests, it is alternatively suggested to use multiple contrast tests vs. total mean and to perform multiplicity adjustment by object merging in the interplay between the R-packages *emmeans* and *multcomp*. This new approach, denotes as *multipleANOM*, allows not only to interpret global factor effects but also local effects between factor levels as adjusted p-values or simultaneous confidence intervals for selected effect measures in generalized linear models. R-code is provided by means of selected data examples.

## 1 The problem

Compared to widely used standard ANOVA, controlling the familywise error rate (FWER) for various multiple factors (and their interactions (IA)) limits its high false positive rate [CvRM^+^16], especially in exploratory analysis [Rub]. Commonly, the easy-to-perform Bonferroni adjustment was proposed [CvRM^+^16], e.g. available in the R-CRAN *library(afex)* [afe]. Furthermore, the standard ANOVA is limited per definition to the interpretation of global effects, whereas one is actually more interested in the individual inference between the factor levels (and their IA) (as long as one does not use simple 2-by-2 designs). Notice, the intention to include more than 2 levels in a design is then also the desire for individual statements, not just one global one. This limited interpretation can be overcome by the similarity of the sum of the squared deviation to the overall mean (as numerator of the F-test) with the maximum of the linear deviations to the overall mean [KBBH13]. I.e. one replaces the standard ANOVA with multiple contrast tests in comparison to the overall mean [PH16], denotes as analysis of means (ANOM).

Furthermore, one replaces the multiplicity adjustment according to Bonferroni by that between multiple models (linear models (lm) or generalized linear models (glm)), taking the correlation between the many contrasts into account. A major advantage of this new approach is the availability of adjusted p-values and compatible simultaneous confidence intervals (for single-step approaches) as well as the applicability of the same principle for numerous effect sizes (such as difference of means, odds ratios, etc.) and models. Another advantage is the possibility of one-sided tests, since the ANOVA F-test is inherently two-sided and thus already conservative. This approach is presented rather formula-less with R-commands by means of example data.

## 2 The multipleANOM approach

ANOVA per se is a wide framework from an one-way layout to high factorial (incomplete) designs, from fixed to mixed effects, from factors with exactly 2 levels to multiple levels, from analyses with or without adjustment against covariates, univariate or multivariate designs, and so on. Here we consider a generally unbalanced design with a few factors (i.e., K=1,2,3), where at least one fixed factor of these has some few levels (J=3,4,5,…,)). Interaction between the factors *ϵ*_*jki*_ are considered and approximate normally distributed errors are assumed (which, however, allow heteroscedasticity): *y*_*jki*_ = *µ*+*ξ*_*j*_ +*ψ*_*k*_ +(*ξψ*)_*jk*_ +*ϵ*_*jki*_ with 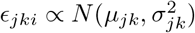

*The first model assumption* is to control for FWER by multiplicity-adjusting the marginal p-values of the ANOVA F-tests [CvRM^+^16].

*The second model assumption* is the hypotheses and power similarity of the sum of the squared deviation to the overall mean in the numerator of the F-test: 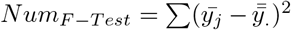 with the effect size of the maximum of the linear contrasts, denoted as analysis of means vs. overall mean (ANOM)[PH16] 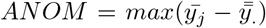, see details in [KBBH13]). I.e. the *multipleANOAM* approach has about the same power compared to ANOVA-whereby there could be tiny power advantages or disadvantages depending on the data and design. The disadvantages should be tolerable, however, in comparison to the more diverse interpretability of both global and local effects.

*The third model assumption* is that the multiplicity adjustment is performed by object combination of the basic models for factor A, factor B and factor AB by the interplay of the *library(emmeans)* [Len20] and the *library(multcomp)* (see the R-code below). The statistical background is the maxT-test which follows a multivariate t-distribution with a correlation matrix R, depending on *n*_*i*_ and the contrast coefficients [Hot16a].

*The fourth model assumption* is modeling of variance heterogeneity by sandwich estimator [Zei06] as a standard approach [HH12]. Variance heterogeneity may be a serious source of bias in any multiple comparison procedure [HH08].

*The fifth model assumption* is the possibility of one-sided inference within the *multipleANOM* approach, whereas ANOVA F-tests are two-sided inherently. Its Bonferroni threshold with *α/*2 is remarkable.

The formal representation of *multipleANOM* is based on R-code, where its object-oriented property is central, e.g. for a two-way layout (with endpoint *y*_*jk*_, two factors f1,f2 and their interaction f1f2):

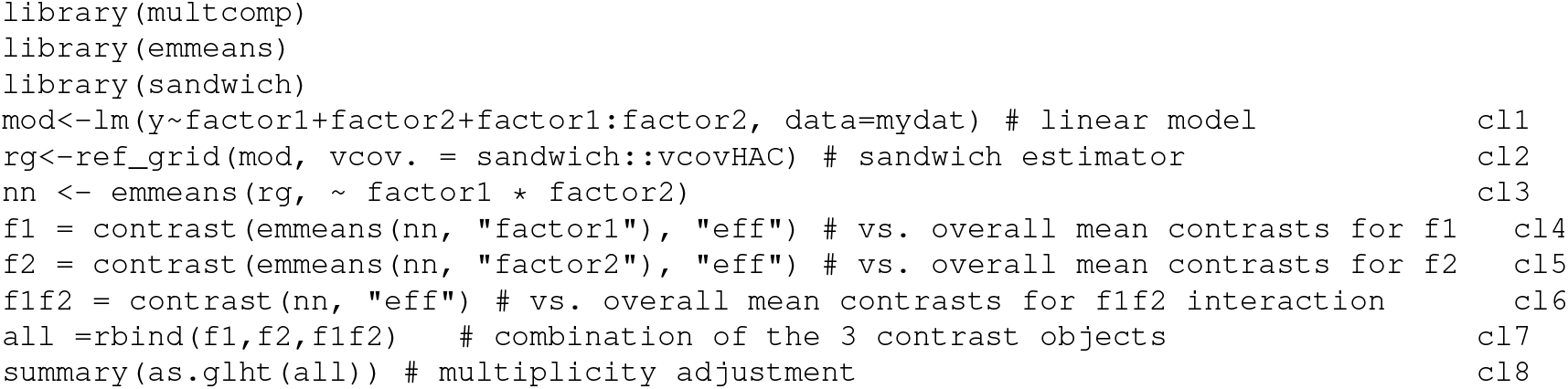

Code line 1 (cl1) shows the usual linear model with interaction term, cl2 the use of the sandwich estimator, and cl3 the estimated marginal means estimators for the factor interaction model. Code lines 4-6 show the contrast estimation for factor 1, 2 and its interaction for the multiple contrast vs. the overall mean. Code line 7 shows the combination of these three objects for simultaneous testing in cl 8.

**In summary**, the *multipleANOM* approach provides adjusted p-values or simultaneous confidence intervals to interpret the global factor effect (by means of *min*(*p*_*ς*_) approach, where *ς* contrasts are included in the particular factor inference as well the individual inference of the factor levels vs. overall mean. What is striking about first use is the large number of individual comparisons compared to the few in ANOVA. I.e., there is not only a hidden multiplicity issue but also a hidden inference issue. This large number can be significantly reduced by *min*(*p*_*ς*_) from ANOVA point of view. Above all, the many individual comparisons are not a disadvantage - on the contrary, the advantage of the simultaneous interpretation of global and local effects. Hereby the contradiction between a-priori and post hoc testing can be overcome in principle (see e.g. [Hot16]). For a long time there has been a curious discussion about post-hoc tests, i.e. multiple comparisons between factor levels if the global ANOVA F-test was previously significant. First, such conditional tests are not conflict-free [KH20], and second, the global F-test for a factor are not of interest at all, neither if it is significant and a post-hoc comparison is indicated (only those are of interest), nor if it is non-significant (Remember: *’absence of evidence is no evidence of absence’* [AB95]). In this context, one should omit the F-tests completely and limit the ANOVA technique to estimate the expected values, variance, and degree of freedom: precisely the estimates of a linear model for qualitative factors.

Some researchers intuitively understand the problems of hidden multiplicity in evaluating factorial design and evaluate the data directly in the cell means model as a pseudo one-way layout, such as the influence of wine varieties (gruener veltliner, zweigelt, pinot noir) and locations (Lower Austria, Burgenland) on wine acidity [PBH^+^21].

Another advantage is the homogeneous use of this *multipleANOM* approach for different effect sizes and model variants based on glm’s [HBW08].

## 3 Case studies

Although one-way layout is not the primary focus of this approach, two case studies are discussed first: a simple one-way layout and a design with a covariate, because already here the advantage of *multipleANOM* over the F-test of ANOVA can be demonstrated. More illustrative is the third example representing a two-way layout with mild interaction.

### 3.1 One-way layout with normal distributed homogeneous or heterogeneous variances

The fatty acid content for different of Bacillus simplex ecotypes are considered [SBKT08] where this unbalanced design reveals heterogeneous variances:

**Figure 1:**
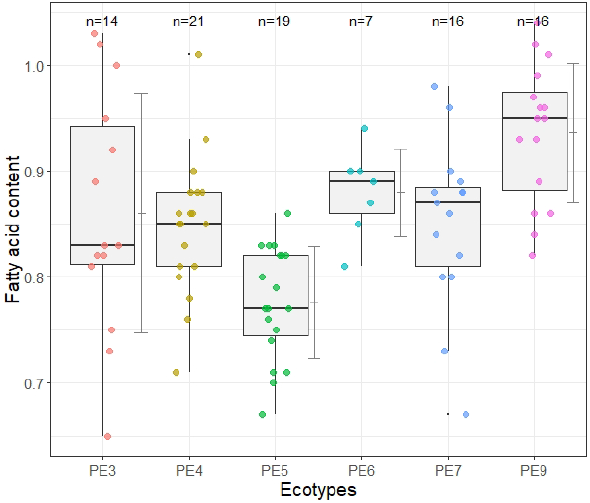
Boxplot fatty acid example: one-way layout

Table 1 demonstrate a p-value of the standard ANOVA which is one order of magnitude lower than that of the new *multipleANOM* approach, but only as long as variance homogeneity is assumed. In this more realistic model, the relation is exactly reversed, and the new approach has a p-value that is two orders of magnitude smaller. This empirical finding impressively shows how important the appropriate modeling of possible variance heterogeneity is. Not surprisingly, all-pair comparisons can show even lower p-values, while the Scheffe test (which adjusts for all possible contrasts) performs quite poorly. In summary, the new approach can make the following statements: i) the factor ecotypes has a significant influence, and ii) species PE5 showing a significant decrease and species PE9 showing a significant increase over the overall mean.

**Table 1:** One-way design: comparing of different approaches

| Method | Variances | Contrast | test stat. t | p-value |
| --- | --- | --- | --- | --- |
| ANOVA | homo | F-test | (8.3) | **0.0000018** |
|  | hetero | F-test | (12.5) | **0.0000066** |
| OverallMean | homo | PE3-GM | 0.4 | 0.9973814 |
|  |  | PE4-GM | -0.4 | 0.9982357 |
|  | min(p) | PE5-GM | -5.1 | **0.0000148** |
|  |  | PE6-GM | 1.0 | 0.8700316 |
|  |  | PE7-GM | -0.2 | 0.9999709 |
|  |  | PE9-GM | 4.9 | 0.0000225 |
|  | hetero, min(p) only | PE5-GM | -6.2 | **0.000000093** |
| All-pairs Tukey | homo, min(p) only | PE9-PE5 | 5.2 | **0.000000089** |
| All-pairs Tukey | hetero, min(p) only | PE9-PE5 | 5.2 | **0.0000000019** |
| All-pairs Bonferroni | homo, min(p) only | PE9-PE5 | 5.2 | **0.00000014** |
| All-pairs Bonferroni | hetero, min(p) only | PE9-PE5 | 5.2 | **0.0000000065** |
| Scheffe | homo, min(p) only | PE9-PE5 | 5.2 | **0.0000027** |

### 3.2 One-way layout with multiple covariates

In reproductive toxicology studies, the weight of pups depends not only on the dose factor of interest, but possibly also on the number of pups within the litter and the duration of gestation [Wes97]. Therefore, analysis of covariance (KOVAR) is indicated i.e., an extension of the one-way ANOVA in the linear model:

**Figure 2:**
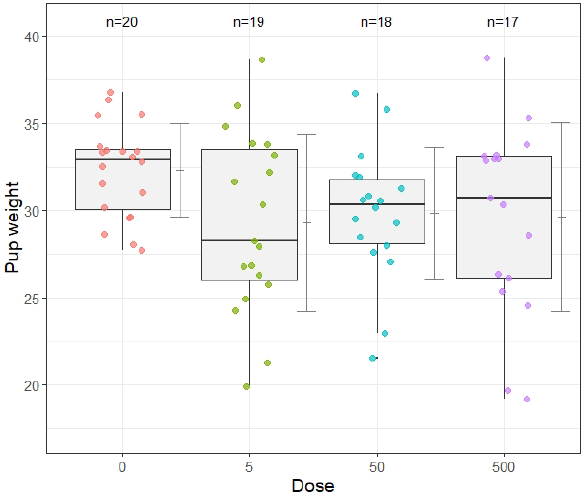
Boxplot litter data

Table 2 demonstrate first inappropriateness of the simple ANOVA approach (ignoring the covariates) with respect to the analysis of covariance (KOVAR) approach. Very clear is the difference between KOVAR and *multipleANOM* (with adjustment vs. covariates): the former is not even significant, while the latter is already significant, which is especially pronounced when variance heterogeneity is modeled. By an order of magnitude lower p-values are obtained if one leaves the pre-testing and immediately uses the Dunnett or even the Williams contrast (and assumption of order restriction), especially with the onesided tests indicated here for the plateau-shaped profile (since only a weight retardation is toxicological relevant). This example clearly shows the fatally qualitatively wrong conclusion of harmlessness, if an unsuitable statistical model is used.

**Table 2:**
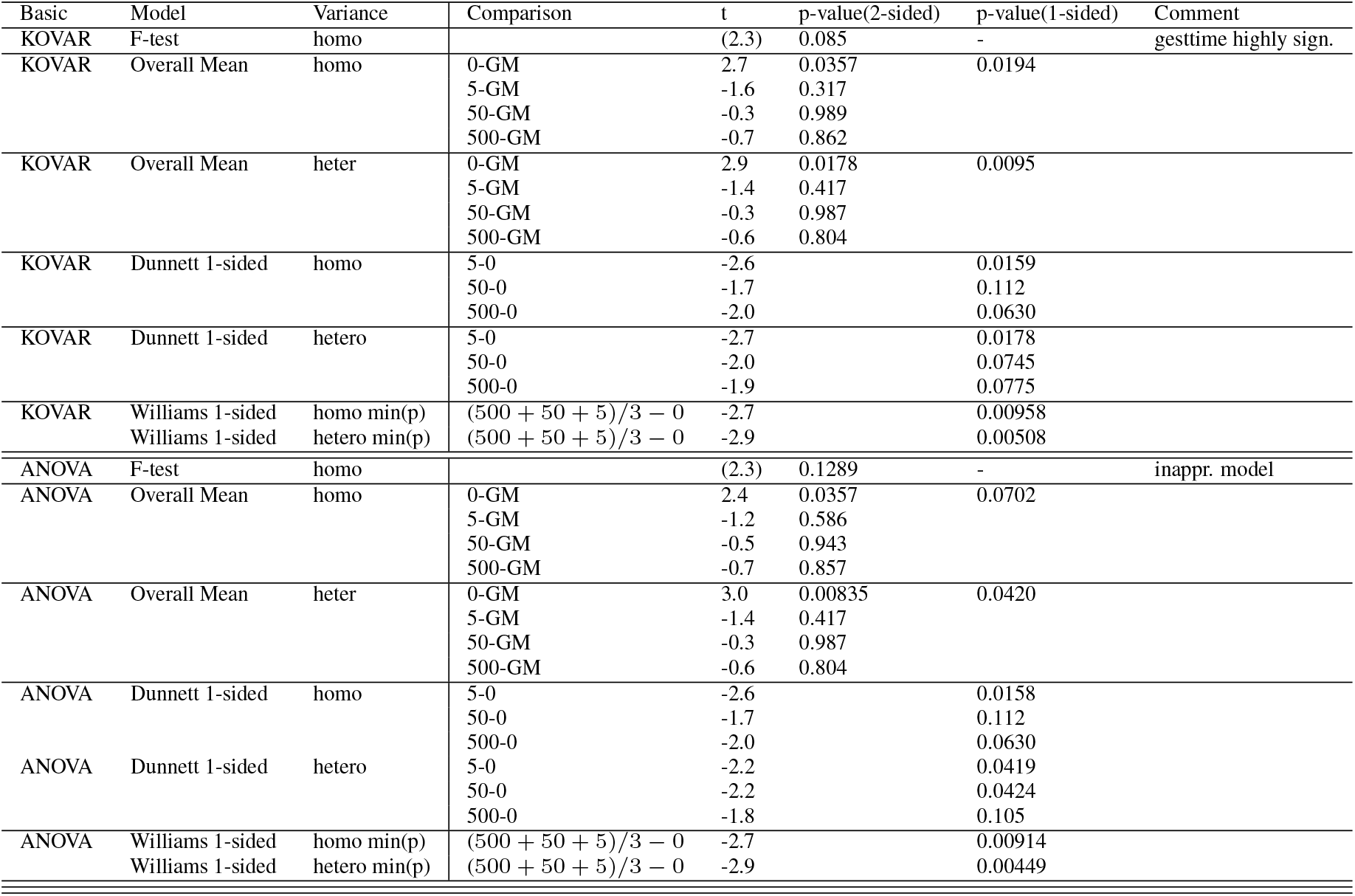
One-way design with multiple covariates: comparing of different approaches

| Basic | Model | Variance | Comparison | t | p-value(2-sided) | p-value(1-sided) | Comment |
| --- | --- | --- | --- | --- | --- | --- | --- |
| KOVAR | F-test | homo |  | (2.3) | 0.085 | - | gesttime highly sign. |
| KOVAR | Overall Mean | homo | 0-GM | 2.7 | 0.0357 | 0.0194 |  |
|  |  |  | 5-GM | -1.6 | 0.317 |  |  |
|  |  |  | 50-GM | -0.3 | 0.989 |  |  |
|  |  |  | 500-GM | -0.7 | 0.862 |  |  |
| KOVAR | Overall Mean | heter | 0-GM | 2.9 | 0.0178 | 0.0095 |  |
|  |  |  | 5-GM | -1.4 | 0.417 |  |  |
|  |  |  | 50-GM | -0.3 | 0.987 |  |  |
|  |  |  | 500-GM | -0.6 | 0.804 |  |  |
| KOVAR | Dunnett 1-sided | homo | 5-0 | -2.6 |  | 0.0159 |  |
|  |  |  | 50-0 | -1.7 |  | 0.112 |  |
|  |  |  | 500-0 | -2.0 |  | 0.0630 |  |
| KOVAR | Dunnett 1-sided | hetero | 5-0 | -2.7 |  | 0.0178 |  |
|  |  |  | 50-0 | -2.0 |  | 0.0745 |  |
|  |  |  | 500-0 | -1.9 |  | 0.0775 |  |
| KOVAR | Williams 1-sided | homo min(p) | $(500 + 50 + 5)/3 - 0$ | -2.7 | | 0.00958 | |
| | Williams 1-sided | hetero min(p) | $(500 + 50 + 5)/3 - 0$ | -2.9 | | 0.00508 | |
| ANOVA | F-test | homo |  | (2.3) | 0.1289 | - | inappr. model |
| ANOVA | Overall Mean | homo | 0-GM | 2.4 | 0.0357 | 0.0702 |  |
|  |  |  | 5-GM | -1.2 | 0.586 |  |  |
|  |  |  | 50-GM | -0.5 | 0.943 |  |  |
|  |  |  | 500-GM | -0.7 | 0.857 |  |  |
| ANOVA | Overall Mean | heter | 0-GM | 3.0 | 0.00835 | 0.0420 |  |
|  |  |  | 5-GM | -1.4 | 0.417 |  |  |
|  |  |  | 50-GM | -0.3 | 0.987 |  |  |
|  |  |  | 500-GM | -0.6 | 0.804 |  |  |
| ANOVA | Dunnett 1-sided | homo | 5-0 | -2.6 |  | 0.0158 |  |
|  |  |  | 50-0 | -1.7 |  | 0.112 |  |
|  |  |  | 500-0 | -2.0 |  | 0.0630 |  |
| ANOVA | Dunnett 1-sided | hetero | 5-0 | -2.2 |  | 0.0419 |  |
|  |  |  | 50-0 | -2.2 |  | 0.0424 |  |
|  |  |  | 500-0 | -1.8 |  | 0.105 |  |
| ANOVA | Williams 1-sided | homo min(p) | $(500 + 50 + 5)/3 - 0$ | -2.7 | | 0.00914 | |
| | Williams 1-sided | hetero min(p) | $(500 + 50 + 5)/3 - 0$ | -2.9 | | 0.00449 | |

### 3.3 Two-way layout with possible interaction

An unbalanced, small *n*_*i*_ two-way design was selected where the litter weight is modeled for the two factors: i) mothers genotype and ii) litters genotype (each with four different genotype levels: mother genotypes [*A, B, I, J*], litter genotype [*a, b, i, j*]):

**Figure 3:**
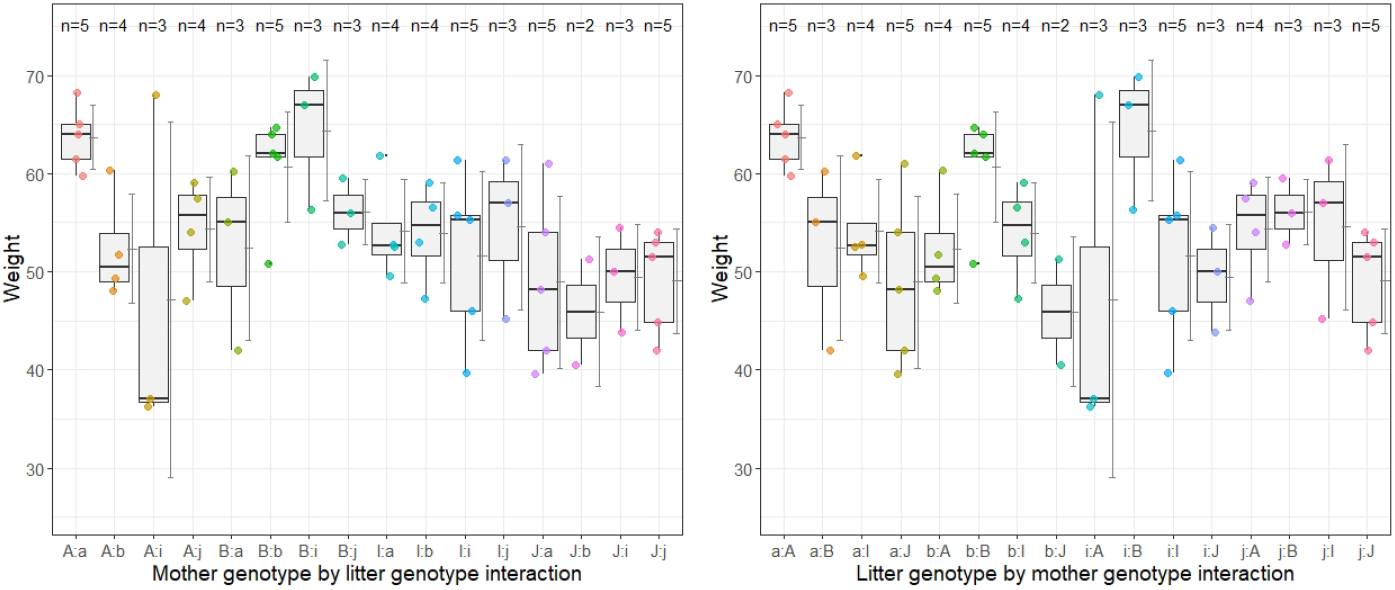
Two-way layout interaction plots

Again, modeling possible variance heterogeneity (or not) is essential to the conclusions especially in this unbalanced, small *n*_*i*_ design. The ANOVA (with variance heterogeneity) would conclude with the results (see Table 3): no effect of the mother genotype, no effect of the litter genotype and no interaction (with homogeneity assumption: the factor litter genotype alone is significant). The original question is with which mother-litter genotype combination the highest weight is achieved. For this the ANOVA seems to be generally unsuitable. Not surprisingly in an unbalanced design with very small *n*_*i*_ and presumably heterogeneous variances, one recognizes large differences between the traditional variance-homogeneous approach and that using sandwich estimators. Put otherwise here, the traditional approach can be quite biased and should be avoided. Also the inappropriateness of the ANOVA is shown, even a clearly non-significant interaction is detected. The essential result is the highest distance from the overall mean for the combination (mother genotypes A, litter genotype a).

**Table 3:** Two-way design: standard ANOVA vs. multipleANOM approach

| Method | Contrast | p-value homo (Bonferroni) | p-value hetero (Bonferroni) |
| --- | --- | --- | --- |
| ANOVA | F-test motgen | 0.775 (0.999) | 0.934 (0.999) |
|  | F-test litgen | **0.0057 (0.0171)** | **0.0480 (0.144)** |
|  | F-test motgen:litgen IA | 0.120 (0.360) | 0.338 (0.999) |
| multiANOM | litgen: a=0 | 0.999 | 0.998 |
|  | litgen: b=0 | 0.999 | 0.999 |
|  | litgen: i=0 | 0.999 | 0.999 |
|  | litgen: j=0 | 0.999 | 0.999 |
|  | motgen: A=0 | 0.999 | 0.999 |
|  | motgen: B=0 | 0.162 | **0.0149** |
|  | motgen: I=0 | 0.999 | 0.999 |
|  | motgen: J=0 | 0.076 | **0.00037** |
|  | IA motgen:litgen A:a=0 | 0.066 | **0.000001** |
|  | IA motgen:litgen A:b=0 | 0.999 | 0.999 |
|  | IA motgen:litgen A:i=0 | 0.862 | 0.998 |
|  | IA motgen:litgen A:j=0 | 0.999 | 0.999 |
|  | IA motgen:litgen B:a=0 | 0.999 | 0.999 |
|  | IA motgen:litgen B:b=0 | 0.476 | **0.0479** |
|  | IA motgen:litgen B:i=0 | 0.208 | **0.0313** |
|  | IA motgen:litgen B:j=0 | 0.999 | 0.596 |
|  | IA motgen:litgen I:a=0 | 0.999 | 0.999 |
|  | IA motgen:litgen I:b=0 | 0.999 | 0.999 |
|  | IA motgen:litgen I:i=0 | 0.999 | 0.999 |
|  | IA motgen:litgen I:j=0 | 0.999 | 0.996 |
|  | IA motgen:litgen J:a=0 | 0.926 | 0.966 |
|  | IA motgen:litgen J:b=0 | 0.882 | **0.0211** |
|  | IA motgen:litgen J:i=0 | 0.996 | 0.119 |
|  | IA motgen:litgen J:j=0 | 0.937 | **0.0165** |

From the user’s point of view, the first striking feature is the large number of 24 p-values compared to the 3 p-values of the ANOVA. On the one hand, the global *min*(*p*_*ς*_) criterion comes to exactly 3 p-values. On the other hand, these many p-values allow exactly the interesting individualized interpretation.

Of course, this *multipleANOM* approach is limited to a few factors. When using factorial experiments one often observes the mistake to include as many factors as possible in the design and to evaluate them with ANOVA based on global F-tests. The more factors, the more likely interactions occur (and not only 2s but 3s, etc.). This is already globally difficult to interpret, increases the conservatism (according to [CvRM^+^16] Bonferroni-adjustment) massively and reduces the power for the primary factor of interest, since these can then only be analyzed with a greatly reduced *n*_*i*_ at the separate levels of the other factors (with interactions).

## 4 Extensions

### 4.1 Testing main factor contrasts when interaction effect is present

The contrasts between the levels of the primary factor are usually tested separately on the levels of the secondary factor if there is a significant interaction or pooled over its levels if there is no interaction. This approach lacks alone in how to test unbiased for existing or without Interaction. Therefore, one can simultaneously test for the contrasts of the factors pooled and per level of the secondary factor [HH].

### 4.2 Using glm’s instead of simple lm’s

Curiously, almost all publications on ANOVA and multiple comparisons are oriented towards normal distribution models. However, there are studies with proportions, counts, time-to-event endpoints,etc., which can be evaluated with comparable approaches. This is possible with the above approach since it is asymptotically based on glm’s. Below is just a simple two-way layout example with a proportion endpoint (percentage surviving trout eggs) with the factors stream location and weeks after placement [HD98]:

The analysis in Faraway’s book [Far02] ends with the conclusion *’We see that both terms are clearly significant’*. Our analysis (glm with add 2 adjustment [AC00]; see the appendix) shows in Table 4 exactly the same outcome with the *min*(*p*_*ς*_) approach, but in addition also a significant interaction, as well as significantly increased survival rates in locations 1 and 2, in week 4 and particularly the highest rates in all locations in week 4.

**Table 4:** Evaluation of a two-way design with a proportion

| No | comparison | test statistics | p-value |
| --- | --- | --- | --- |
| 1 | 1 location | 7.234776 | 0.0000001 |
| 2 | 2 location | 6.156309 | 0.0000001 |
| 3 | 3 location | 2.461923 | 0.286 |
| 4 | 4 location | 4.082125 | 0.0012 |
| 5 | 5 location | -14.157383 | 0.0000001 |
| 6 | wk11 | -4.966372 | 0.000019 |
| 7 | wk4 | 7.970087 | 0.0000001 |
| 8 | wk7 | -1.548965 | 0.937 |
| 9 | wk8 | -2.449478 | 0.295 |
| 10 | 1 wk11 | 3.781131 | 0.0044 |
| 11 | 2 wk11 | 1.931773 | 0.709 |
| 12 | 3 wk11 | -1.165653 | 0.997 |
| 13 | 4 wk11 | 1.720501 | 0.858 |
| 14 | 5 wk11 | -6.401059 | 0.0000001 |
| 15 | 1 wk4 | 3.591705 | 0.0091 |
| 16 | 2 wk4 | 4.289696 | 0.0005 |
| 17 | 3 wk4 | 4.466290 | 0.0002 |
| 18 | 4 wk4 | 4.072643 | 0.00128 |
| 19 | 5 wk4 | -5.137541 | 0.000008 |
| 20 | 1 wk7 | 3.868902 | 0.0030 |
| 21 | 2 wk7 | 1.916483 | 0.721 |
| 22 | 3 wk7 | -0.140459 | 0.999 |
| 23 | 4 wk7 | 1.051015 | 0.999 |
| 24 | 5 wk7 | -11.032345 | 0.0000001 |
| 25 | 1 wk8 | 2.449776 | 0.294 |
| 26 | 2 wk8 | 2.847300 | 0.107 |
| 27 | 3 wk8 | -0.878667 | 0.999 |
| 28 | 4 wk8 | -2.226895 | 0.460 |
| 29 | 5 wk8 | -9.645250 | 0.0000001 |

## 5 Summary

As an alternative to standard ANOVA, *multipleANOM* is proposed, which directly provides multiplicity-adjusted p-values (or simultaneous confidence intervals) for contrasts compared to the overall mean for the main and interaction effects, and modeling variance heterogeneity. The *multipleANOM* approach allows various extensions, since glm’s are underlying under the limitation to asymptotic tests.

There is no question that adjusting against hidden multiplicity reveals a conservative behavior relative to standard ANOVA. However, in the mostly non-a priori powered studies, some conservatism is preferable to a massive false positive rate. Implicit in this new approach is a restriction of factorial designs to only a few factors. In my opinion, this also applies to the standard ANOVA, since one then has to expect interactions which on the one hand make the interpretability more difficult and on the other hand the power for the inference of the factors of interest (and their levels) is dramatically reduced.

## Acknowledgment

My thanks to Dr. Lenth, University of Iowa, for the information on contrast formulation in the library(emmeans) and their combination for simultaneous testing.

## 6 Appendix

### data(foster)

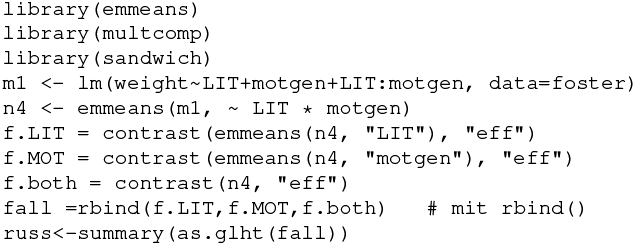

### data(troutegg)

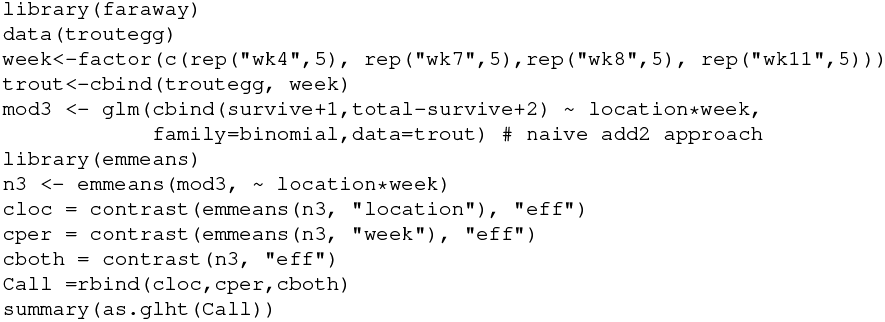

